# Human Urine-derived SIX2-positive renal progenitor cells improve kidney injury in an IRI mouse model

**DOI:** 10.1101/2025.07.02.662781

**Authors:** Claus Kordes, Lucas-Sebastian Spitzhorn, Martina Bohndorf, Audrey Ncube, Chantelle Thimm, Lars Erichsen, Wasco Wruck, James Adjaye

## Abstract

Cells with the characteristics of renal progenitor cells are shed in urine. The aim of this study was to investigate whether these SIX2-positive urine-derived renal progenitor cells (UdRPC) have therapeutic potential in treating or managing acute to chronic kidney injuries. Human UdRPC were obtained from a woman aged 35 years, expanded, and characterized *in vitro* and finally transplanted unilaterally under the renal capsule of mouse kidneys that had undergone ischemia reperfusion injury (IRI). Analyses of blood serum from the mice revealed that the transplanted human UdRPC transiently influenced the secretome of the animals by triggering the release of inflammatory and angiogenic factors. Of obvious therapeutic relevance, however, was the finding that the transplanted cells suppressed renal fibrosis resulting from IRI. The expression of genes linked to inflammation and chronic kidney disease was also reduced in mouse kidneys in the presence of UdRPC. Ultimately, the human UdRPC were able to alleviate the severity of the injury through the amelioration of fibrosis but were unable to restore complete kidney function within 21 days. Nevertheless, the findings of this study demonstrate a significant influence of transplanted UdRPC on kidney fibrosis after acute to chronic injury presumably via an altered secretome.

## INTRODUCTION

Acute kidney injury (AKI) is triggered by ischemia, hypoxia, or nephrotoxicity which initiate inflammatory processes that can lead to a chronic progression of the disease and renal fibrosis ^1,2,3^. Chronic kidney disease (CKD) is further initiated independently from AKI by diabetes or hypertension but can also be caused by the natural ageing processes ^4,5^. The prevalence of CKD is increasing worldwide, and it has been estimated to represent the seventh leading cause of death worldwide ^6^. In Europe, CKD affects one in 10 people and is therefore a serious problem for the healthcare system ^4^. Over the years, reports have described urine as a promising source for stem cells in clinical use ^7^. Urine has been shown to harbor urine-derived renal progenitor cells (UdRPC). UdRPC express key kidney transcription factors such as SIX2 ^8,9^. Furthermore, their identity as kidney progenitor cells which can differentiate into podocytes has been reported ^10^. Their capacity to differentiate into renal tubule cells can be assumed because the UdRPC express SIX2. This transcription factor regulates self-renewal of bipotent nephron progenitor cells during kidney organogenesis ^11^. Apart from their kidney specific features, UdRPC expanded *in vitro* on plastic exhibit properties of mesenchymal stem cells such as a trilineage differentiation into adipocytes, osteocytes, and chondrocytes as well as secretion of immunomodulatory cytokines and angiogenic growth factors ^9^. First attempts have been made by several groups to estimate the therapeutic value of transplanted urine stem cells using rodent models of AKI and CKD with beneficial effects on kidney function and regression of renal fibrosis ^12,13,14,15,16^. However, mechanisms initiated by SIX2-expressing kidney progenitor cells that contribute to the alleviation of renal fibrosis remain largely unknown.

To explore this, AKI was performed in the present study by transient, unilateral clipping of the renal pedicle to initiate ischemia reperfusion injury (IRI) in mice. IRI interrupts blood circulation of the kidneys resulting in endothelial cell damage, tubular cell injury, and inflammation which can trigger fibrosis ^17^. Depending on the duration of the ischemic status, the consequences of AKI can vary between mild as well as severe forms with progression into CKD as has been shown for the well-established model organism mouse ^18,19^. Unilateral transient clipping of the renal pedicle of mice for 45 min as performed in the present study induced AKI to CKD accompanied by renal fibrosis and impaired kidney function. Injection of human UdRPC under the capsule of IRI-injured kidneys, however, was unable to completely restore kidney function but prevented the progression of renal fibrosis as reported in this study.

## RESULTS

### Characterization and transplantation of UdRPC

Isolated human UdRPC showing a spindle-shaped cell morphology expressed CD133 and the kidney progenitor markers SIX homeobox 2 (SIX2) and Cbp/p300 interacting transactivator with Glu/Asp rich carboxy-terminal domain 1 (CITED) (Fig. 1A). Flow cytometry analysis revealed that UdRPC express the well-established kidney progenitor cell surface proteins CD24 (99%), CD106 (58%), and CD133 (99%) (Fig. 1B). UdRPC could be differentiated into both podocytes expressing NPHS1 adhesion molecule (NPHS1/nephrin), α-actinin 4, synaptopodin (SYNPO), and NPHS2 stomatin family member (NPHS1/podocin) (Fig. 1C) as well as tubular cells expressing Na^+^/K^+^-ATPase, CLCNKB (chloride voltage-gated channel Kb), SOX17 (SRY-box transcription factor 17), and ZO-1 (zonular occludens-1) (Fig. 1D). Approximately 1x10^6^ UdRPC in 20 µl PBS were injected under the capsule of the left kidney of sham operated mice (group 1), while the vehicle PBS was injected in the same way into the contralateral kidney (Fig. 2A). The left kidney of group 2 and 3 underwent IRI and animals of the IRI group 2 received PBS injections in both kidneys, while mice of the IRI group 3 received 1x10^6^ UdRPC into the clipped left kidney and only PBS was injected into their contralateral kidney (Fig. 2A).

**Figure 1:**
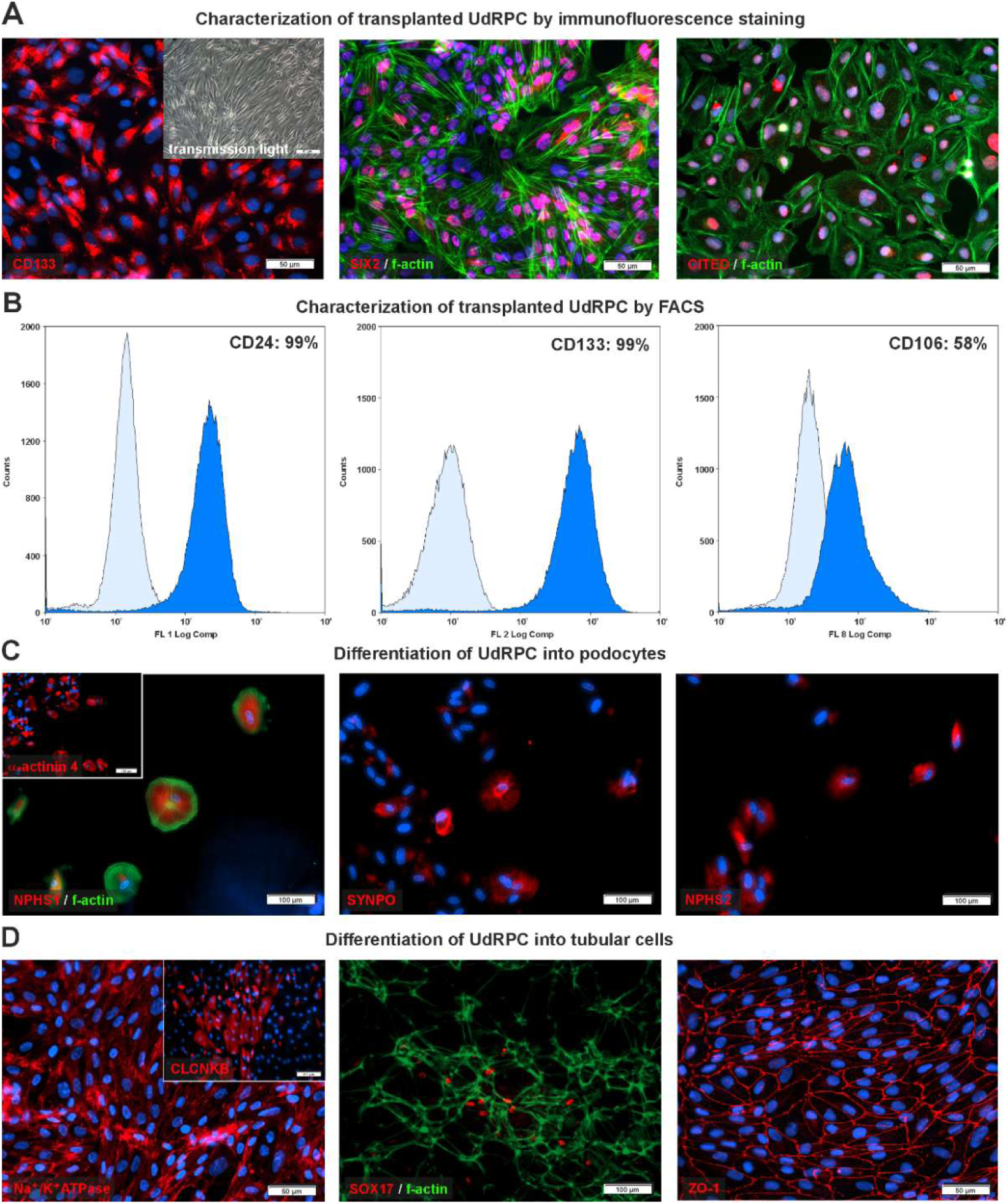
Characterization of UdRPC by immunofluorescence and flow cytometry. (**A**) Transmission light microscopic image of UdRPC showing spindle-shaped cell morphology (insert). Immunofluorescence staining of UdRPC using antibodies against the kidney progenitor cell markers CD133, SIX2, and CITED (red). The cell nuclei were stained with Hoechst 33258 (blue) and the actin cytoskeleton was labelled by phalloidin (green). Scale bars represent 50 µm. (**B**) The renal progenitor cell markers CD24, CD106 (VCAM1), and CD133 were detectable as cell surface proteins on UdRPC by flow cytometry. Dark blue histograms indicate the fluorescence-labelled antibodies against CD24, CD106, or CD133, respectively, whereas light blue histograms represent antibody isotype controls. (**C**) Podocytes differentiated from UdRPC expressed characteristic podocyte markers NPHS1, β-actinin 4 (insert), SYNPO, and NPHS2 (red). The cytoskeletal marker phalloidin is shown in green. (**D**) UdRPC differentiated into tubular cells showed tubular-associated markers Na^+^/K^+^-ATPase, CLCNKB (insert), SOX17, and ZO-1 (red) at protein level, along with phalloidin. Scale bars indicate 50 and 100 μm.

**Figure 2:**
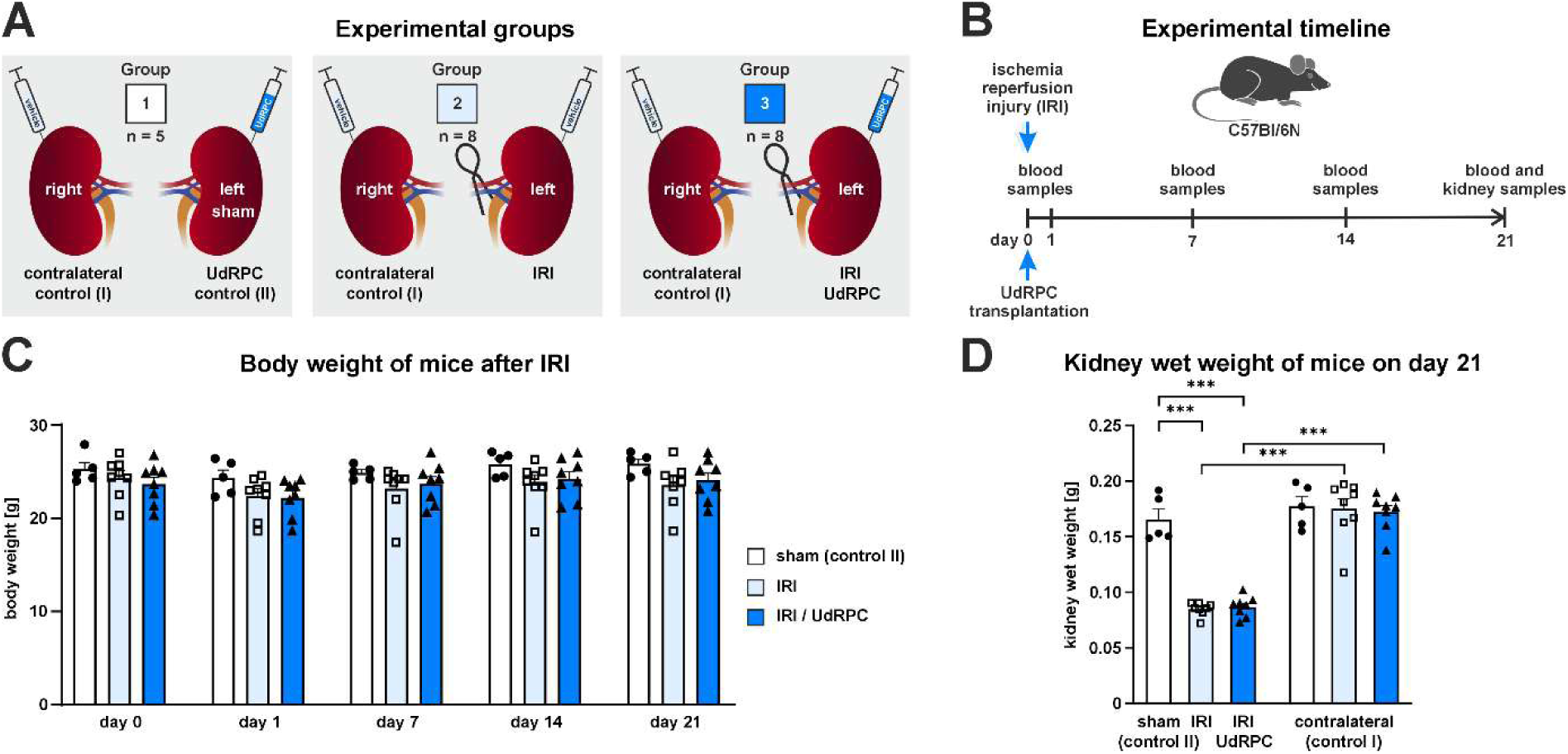
Experimental procedure and health status of mice as indicated by body and kidney weight. (**A**). Group 1 was sham operated and received UdRPC injection into the left kidney. IRI-induced AKI was triggered by unilateral transient clipping of the left renal pedicle in group 2 and 3. The injured left kidney of the group 3 received an injection of UdRPC whereas only the vehicle PBS was injected into the left kidney of group 2. The contralateral (right) kidneys always received the vehicle in all groups. (**B**) Timeline of the experimental procedure. (**C**) Body weight of mice was determined prior to surgery (day 0) and on day 1, 7, 14, and 21. (**D**) Kidney wet weight on day 21. Data are indicated as mean ± SEM. Statistical significance was tested by one way ANOVA. Significant differences were ***p<0.001.

As indicated by the experimental timeline (Fig. 2B), blood samples were taken and the body weights were monitored prior to surgical intervention at day 0 as well as after surgery at day 1, 7, 14, and 21. The body weight of mice remained unchanged during the course of the experiment which implies that the surgery had no significant influence on the health status of the mice (Fig. 2C; supplemental Table S2). After 21 days, the animals were sacrificed, and the kidney wet weight was determined. IRI triggered a significant weight loss in the left kidney of mice in groups 2 and 3 in comparison to the contralateral kidneys (control I) and the left sham treated controls (control II) (Fig. 2D). No significant differences were observed in the kidney wet weight between the group 2 (IRI) and 3 (IRI/UdRPC) (Fig. 2D).

### Renal function after IRI

The concentration of blood urea nitrogen (BUN) in the serum of mice was increased in both IRI groups (group 2: 13 mmol/L; group 3: 10 mmol/L) on day 1 after surgery but only the BUN levels of the IRI group 2 reached significance when compared to the sham treatment (group 1: 8 mmol/L) (Fig. 3A). A significant increase in BUN levels was also observed for the IRI group 2 on day 7 post-surgery when compared to the sham control (group 1), whereas BUN levels of the group 3 (IRI/UdRPC) were elevated but not significantly different from the sham control throughout the entire experiment (Fig. 3A), suggesting better recovery of the kidneys after AKI in the presence of UdRPC. A similar pattern emerged for the creatinine levels in the blood serum. Serum creatinine increased in both IRI groups up to approximately 30 µmol/L on day 1 post-surgery compared to baseline (day 0) and the sham control (day 1), which showed a concentration of approximately 24 µmol/L (Fig. 3B). On day 7 post injury, creatinine levels of the IRI group 2 were still significantly increased, whereas in the IRI group 3 with transplanted UdRPC no significant differences compared to baseline and sham control was observed (Fig. 3B), thus indicating supportive effects of UdRPC on injured kidney tissue.

**Figure 3:**
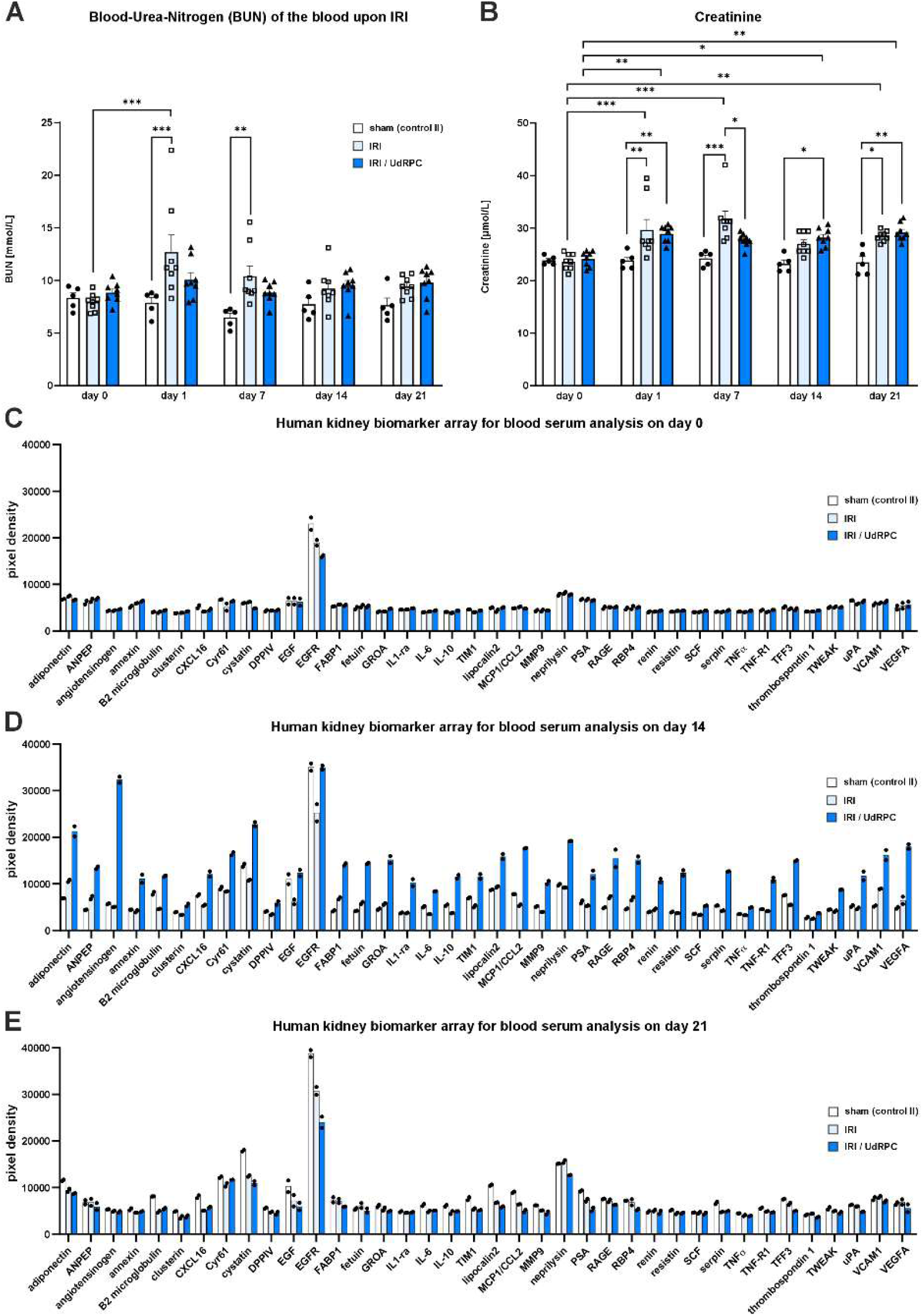
Blood serum analyses. (**A**) Blood-urea-nitrogen (BUN) and (**B**) creatinine concentrations as indicators of kidney injury were measured in the blood serum prior to surgery (day 0) and on day 1, 7, 14, and 21. Data are shown as mean ± SEM (n = 5-8). Statistical significance was tested by one way ANOVA. Significant differences were *p<0.05, **p<0.01, ***p<0.001. (**C-E**) Proteome Profiler Human Kidney Biomarker Arrays have been used to monitor cytokines and growth factors associated with kidney diseases in the blood serum of mice at day 0, 14, and 21. The serum samples were pooled from different animals according to their experimental group assignment to receive sufficient protein amounts. The bars indicate mean values, and the dots represent individual values of double measurements.

### Proteome analyses

To gain insight into the underlying mechanisms of transplanted UdRPC on kidney function, the blood serum of the mice was analyzed prior to surgery (day 0) and on day 14 and 21 after AKI using proteome analyses that include kidney biomarkers (Fig. 3C-E). Proteins with an influence on vascular system were found to be increased in the blood serum of the IRI/UdRPC group 3, particularly on day 14 (Fig. 3D). The renin/angiotensin system capable of elevating blood pressure was more pronounced on this day in the UdRPC transplantation group 3. Moreover, vascular endothelial growth factor A (VEGFA), trefoil factor 3 (TFF3), and urokinase (uPA) are factors that promote the survival and proliferation of vascular cells or facilitate thrombolysis, were more abundant in group 3. In addition, adipokines such as resistin, adiponectin, and serpine are known to exert effects on endothelial cells and their expression was found to be elevated in IRI/UdRPC group 3 at day 14 (Fig. 3D). On the other hand, an increased release of factors such as interleukine-6 (IL-6), C-C motif chemokine ligand 2 (CCL2/MCP1, monocyte chemoattractant protein 1) and tumor necrosis factor α (TNFα), which are associated with inflammatory processes and elevated in injured tissue, was found in the IRI/UdRPC transplant group (Fig. 3D). In contrast, the increased expression of the immunosuppressive factors interleukine-10 (IL- 10) and interleukin-1 receptor antagonist (IL-1ra) as well as the extracellular matrix (ECM)-degrading matrix metalloproteinase 9 (MMP9) implies potential beneficial effects of UdRPC on injured kidneys (Fig. 3D). Unlike the situation on day 14, the serum proteome analysis on day 0 and 21 showed minor differences between the different experimental groups (Fig. 3C, E; supplemental Fig. S1A-C), suggesting that UdRPC transiently influenced receiving kidneys at an early stage after transplantation as already indicated by the serum BUN and creatinine analyses (Fig. 3A, B).

Cytokines released by the UdRPC into the culture medium were investigated by a proteome array covering 105 human cytokines to assess the origin of cytokines in the blood of mice on day 14 after transplantation of UdRPC (Fig. 4A-D; supplemental Fig. S2). Individual factors such as colony stimulating factor 2 (CSF2/GMCSF), growth and differentiation factor (GDF15), insulin-like growth factor binding proteins 2/3 (IGFBP2/3), and VEGFA emerged in particular due to their quantity (Fig. 4A). The cytokines released by cultured UdRPC were subjected to GO term analysis with regard to biological processes and cellular components. It was of interest that among the biological processes “tissue homeostasis” and “wound healing” were found (Fig. 4B; supplemental Table S4), which indicated a support of the injured kidney tissue by the detected cytokines. A comparison with the cytokines released in the blood serum on day 14 suggested that the factors found in the blood of the mice of the IRI/UdRPC group 3, such as IL-1ra, IL-10, MMP9, and TNFα, apparently originated from mouse cells, as these were not clearly evident in the UdRPC culture medium (Fig. 4D). In contrast, receptor for advanced glycosylation end product-specific (RAGE) could be distinguished as a factor associated with IRI in mouse blood. It should be emphasized that VEGFA as part of the biological processes “tissue homeostasis” and “wound healing” (Fig. 4B) and the majority of the GO terms referring to cellular components (Fig. 4C; supplemental Table S4) was clearly prominent under all conditions (Fig. 4D).

**Figure 4:**
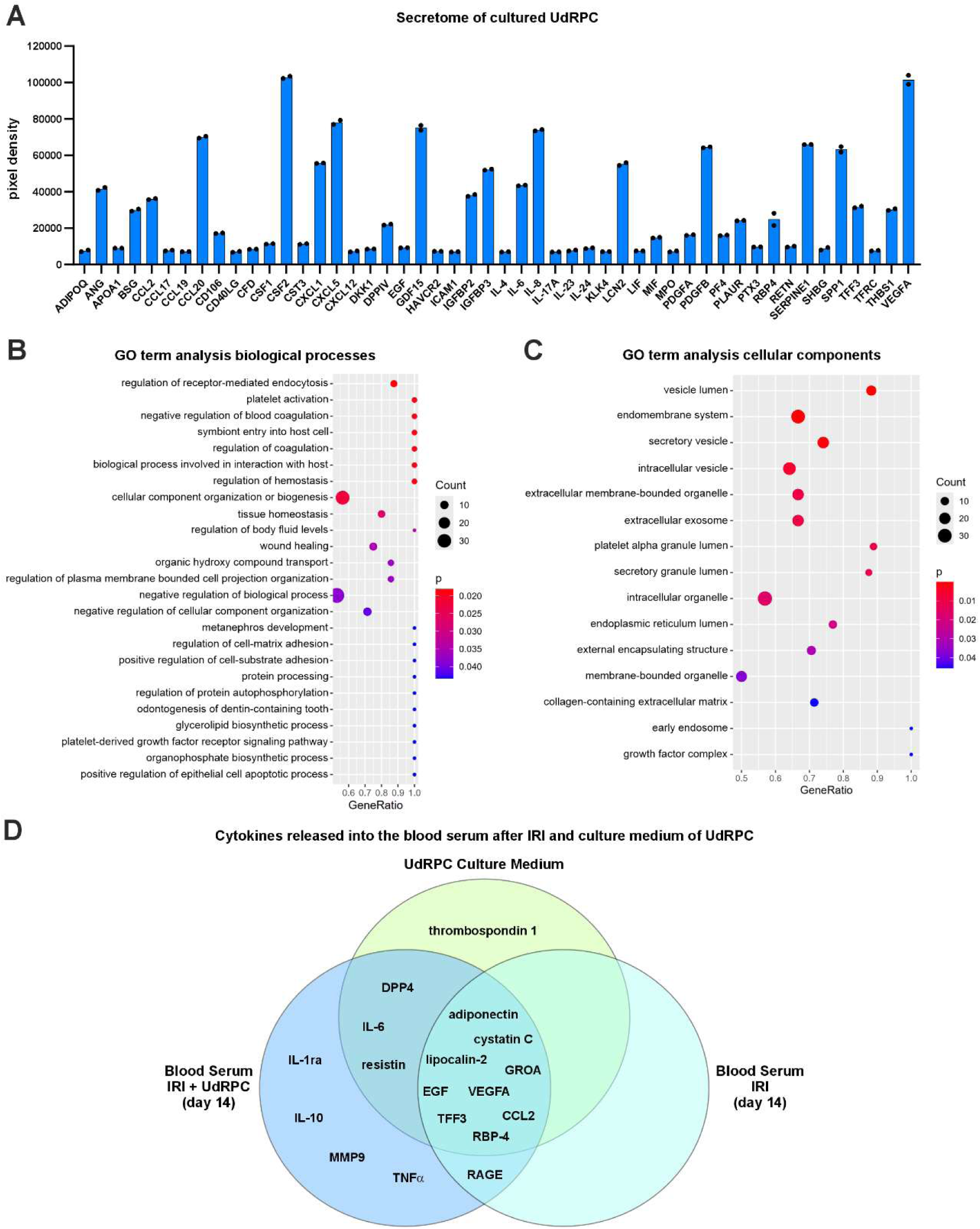
Secretome analysis of UdRPC. (**A**) Cytokines released by UdRPC into the culture medium were analyzed by the Proteome Profiler Human XL Cytokine Array covering 105 cytokines. Only the expressed cytokines that have a detection-p-value below a threshold of 0.05 are shown. The bars indicate mean values, and the dots represent individual values of double measurements. GO over-representation analysis was performed to identify gene ontology (GO) terms of (**B**) biological processes and (**C**) cellular components associated with the cytokines released by UdRPC as indicated by the aforementioned proteome array. (**D**) Matching cytokines from the Proteome Profiler Human Kidney Biomarker Array Kit (ARY019), which was used to analyze the blood serum of the mice 14 days after IRI, and the Proteome Profiler Human XL Cytokine Array Kit (ARY022B), which was applied to identify the cytokines released by UdRPC into the culture medium, were compared in a Venn diagram. In addition to the UdRPC culture medium, the blood sera of mice after IRI were illustrated with and without transplanted UdRPC for day 14 after kidney injury.

### Analyses of kidney fibrosis

Matrix degrading enzymes like MMP9 detected in blood serum proteome analysis prompted us to investigate the ECM deposition of the injured mouse kidneys by Sirius Red staining in response to transplanted UdRPC. The animals from both control groups (control I: contralateral kidney; control II: sham treatment with UdRPC) were combined to obtain a robust mean value as a reference value to analyze fibrosis. This was justified because the kidneys of the animals in both controls showed no differences in kidney weight or the molecular biology and histology analyses as described in the following. Quantification of the Sirius Red-stained areas revealed a significant increase of matrix protein deposition in mice of the IRI group 2 on day 21 and indicated kidney fibrosis (Fig. 5A, B). In contrast, the transplantation of UdRPC into the injured kidney prevented excessive ECM production (Fig. 5A, B). Furthermore, the expression of fibrosis-associated genes such as connective tissue growth factor (*Ctgf/Ccn2*) remained low despite AKI when UdRPC were transplanted (Fig. 5C). Reduced kidney fibrosis in the IRI group 3 with UdRPC was further manifested by the expression analysis of collagen genes. The expression of the *Col1α2* and *Col3α1* chains was significantly lower in the IRI/UdRPC group 3 than in the IRI group 2 (Fig. 5D, E) whereas the expression of the *Col4α1* chain and *Sox9* only showed a tendency towards reduction after transplantation of UdRPC (Fig. 5F, G). Significantly reduced expression was observed for the inflammation-associated genes *Ccl2* and intercellular adhesion molecule 1 (*Icam1*) (Fig. 5H, I) in the mouse kidneys after IRI and transplantation of UdRPC compared to IRI group 2. This further indicated a lower inflammation of the injured kidney tissue in the presence of UdRPC.

**Figure 5:**
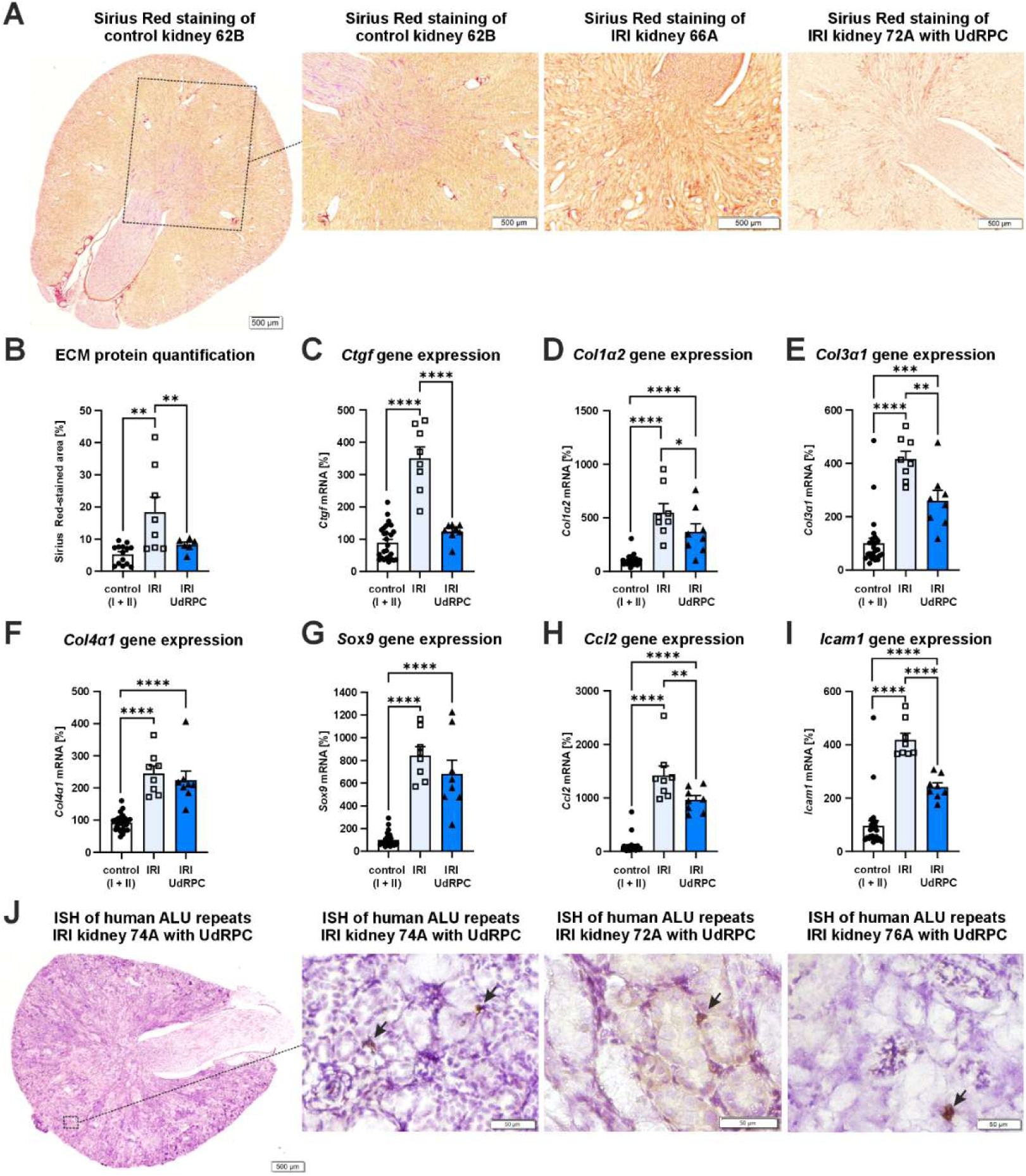
Assessment of kidney fibrosis and detection of transplanted UdRPC. (**A**) Kidney fibrosis was quantified by Sirius Red staining that highlights ECM proteins in red. A whole kidney section scan of a mouse kidney (mouse 62B, control I) stained by Sirius red is shown (left). A detailed enlargement of this section as an example for images used for quantification of Sirius Red-stained areas is also provided. To the right are examples of kidney tissues with significant fibrosis on day 21 after IRI (mouse 66A) and IRI with transplanted UdRPC (mouse 72A) are shown. The scale bars indicate 500 µm. (**B**) Quantification of Sirius Red intensities was performed using images shown above. The values of contralateral (control I) and sham (control II) tissues have been combined and compared with Sirius Red stained kidney sections of group 2 (IRI) and 3 (IRI/UdRPC). (**C-I**) Expression analyses of the fibrosis-associated genes Ctgf, Col1α2, Col3α1, Col4α1, and Sox9 as well as the inflammation-associated genes Ccl2 and Icam1 in mouse kidney tissues by qPCR. Data are indicated as mean ± SEM (n = 8-26). Statistical significance was tested by one way ANOVA. Significant differences were *p<0.05, **p<0.01, and ****p<0.0001. (**J**) Transplanted human UdRPC were detected by ISH using human-specific ALU repeat probes (brown, black arrows). The cell nuclei were stained with hematoxylin (blue). The image on the left shows an overview of an entire kidney section (mouse 74A, IRI/UdRPC). A detailed enlargement of the same animal (74A) is shown to the right and images of two other mice from the transplantation group after IRI (72A and 76A) exhibiting brown cell nuclei of human cells are also provided. The scale bars represent 50 µm.

### Transplanted human UdRPC in mouse kidney

To detect the presence of human UdRPC in mouse kidneys, ISH of human-specific ALU repeats was performed. The transplanted UdRPC were predominantly found in the renal cortex at low cell numbers and mainly located between the tubules where the capillary system is located as shown for 3 mice that received UdRPC (Fig. 5J). According to our ISH analysis only a small number of UdRPC survived in the kidneys. It can be assumed that most of the human cells were presumably eliminated by the immune system during the 21-day period which provides an explanation for the transiently altered blood serum secretome on day 14 (Fig. 3D). However, the positive effect of transplanted UdRPC on prevention of kidney fibrosis persisted (Fig. 5B).

## DISCUSSION

The aim of this study was to test if urine-derived SIX2-positive UdRPC can support injured kidney and thus establish and confirm their therapeutic value and application. Research by other groups has already indicated this ^12,13,14,15,16^. The present study demonstrated that UdRPC have the ability to differentiate into podocytes and tubular cells, as expected for renal progenitor cells. Thus far, very few attempts were made to unravel the mechanisms underlying positive effects of transplanted urine progenitor cells on injured kidneys. For example, exosomal miRNA species such as miR-146-5p and IL- 10 have already been associated with beneficial effects of urine stem cells ^13,14^. However, it is more likely that there is a broad spectrum of factors released by urine progenitor cells influencing the recovery of injured kidney tissue.

In the present study, quantification of ECM proteins in renal tissue using Sirius Red staining revealed that the transplanted UdRPC significantly prevented the development of renal fibrosis as seen in the IRI model without transplanted cells. A possible mechanistic approach to explain this was provided by secretome analyses in the present study. When the proteome of mouse serum and UdRPC culture supernatant were compared, IL-1ra and IL-10, MMP9, and TNFα were found to be increased only in the serum of mice of the IRI/UdRPC group 3. Presumably, this was a response of the mouse tissue to factors released by the transplanted UdRPC, as these proteins were not clearly detectably released by UdRPC into the culture medium. IL-1ra binds IL-1 and has an anti-inflammatory effect ^20,21^, as observed for IL-10. In line with this, a decrease in inflammation in mouse kidneys with IRI and transplanted UdRPC was indicated by a reduced expression of the genes *Ccl2* and *Icam1*.

VEGFA was among the factors produced by UdRPC that were also prominent in the blood serum of mice of the group 3 (IRI/UdRPC). While VEGFA is well known to stimulate angiogenesis and able to counteract ischemia, PDGFB secreted by endothelial cells attracts pericytes which maintain the vasculature ^22,23,24^. According to our proteome analyses, PDGFB is also released from UdRPC into the cell culture medium and can therefore potentially influence the regenerative process. IGFBP2/3, VEGFA, and GDF15 were also among the factors released by UdRPC. Although IGFBP bind and transport insulin-like growth factors (IGF), IGFBP2/3 seem to promote angiogenesis, too ^25,26^. GDF15, in turn, is apparently secreted by senescent endothelial cells presumably to facilitate blood vessel regrowth ^27,28^. Hence, several factors released by UdRPC have an effect that promotes the vascular system. It is interesting to note that an increase in the density of the peritubular capillary system counteracts CKD and reduces kidney fibrosis ^29^. CSF2 is another factor released by UdRPC. In mice with targeted deletion of CSF2 bleomycin-induced lung fibrosis is more severe than in wild type mice apparently by lowered prostaglandin E2 levels ^30^. This can promote immune cell but also sympathetic nervous system activation which both contribute to fibrogenesis ^31,32^. Thus, immunomodulatory and angiogenic mechanisms triggered by UdRPC can explain their anti-fibrotic effect reported herein. However, it remains to be investigated whether the transplanted UdRPC have actually led to a change in the vascular system. This requires further extensive analyses, which should include the aforementioned cytokines.

## Limitations of the study

The sera of several animals had to be pooled in order to obtain sufficient sample material for the proteome analyses, as only small volumes of blood serum can be obtained from living mice. This means that individual variations between the animals in a group cannot be mapped. Furthermore, the low number of UdRPC still present in the mouse tissue between tubules after 21 days suggests that most of the human cells either did not engraft into the kidneys or were rejected by the immune system, as the immunocompetent animals of the C57Bl/6N strain did not receive immunosuppression. Moreover, the limitation of the observation period to 21 days did not enable investigation of potential beneficial long-term effects of UdRPC on mouse kidneys. It is also not clear whether the immunosuppressive IL- 10 and IL-1ra as potential modulators of inflammatory processes counteracted the effect of other pro-inflammatory cytokines (e.g. TNFα and IL-6) or whether the elevated levels of IL-6 in blood serum even had a favorable effect on the regenerative process via the induction of increased endothelial cell proliferation by triggering VEGFA expression ^33,34^.

## Conclusion

Despite these limitations, the present study has unveiled an anti-fibrotic effect of transplanted UdRPC in ischemia-induced acute to chronic renal injury in mice. Anti-inflammatory and angiogenic factors could have mediated the positive effects of UdRPC on the regenerative processes of the mouse kidneys. Further studies are required to identify individual factors in the secretome of UdRPC that can be used in the prevention or treatment of renal fibrosis in patients.

## MATERIALS AND METHODS

### Characterization of UdRPC

Human UdRPC (UF31) were isolated from the urine of a 35-year-old healthy female donor of African origin. Informed consent was obtained by the donor of the urine cells and ethical approval for their use in research was given by the ethical commission of the Heinrich Heine University. All methods were performed in accordance with the relevant guidelines and regulations. UdRPC were initially selected from urine by their adherent growth on plastic as proposed for MSCs. The proliferating UdRPC were expanded using culture medium composed of 50% Dulbecco’s Modified Eagle Medium (high glucose) and 50% keratinocyte medium supplemented with 5% fetal bovine serum (FBS), 0.5% non-essential amino acids, 0.25% GlutaMAX, 0.5% penicillin/streptomycin, and basic fibroblast growth factor (5 ng/ml, Cat# 100-18B, PreproTech Germany, Hamburg, Germany) at 37°C and 5% oxygen. If not otherwise indicated, all components of the culture medium were purchased from Gibco (Carlsbad, CA, USA). UdRPC were fixed with 4% paraformaldehyde (PFA) and incubated with primary antibodies directed against SIX2 (#H00010736-M01, Abnova, Taipei City, Taiwan), CD133 (#C2514-90B, US Biological; PA2049, Boster Biological), Cbp/p300 interacting transactivator with Glu/Asp rich carboxy-terminal domain 1 (CITED1, #PA5-40585, Thermo Fisher Scientific) as well as corresponding secondary Cy3-, Alexa Fluor 555-, or Alexa Fluor 488-labelled antibodies (Thermo Fisher Scientific) and Hoechst 33258 dye (Merck, Darmstadt, Germany) using a standard protocol. The actin cytoskeleton (f-actin) was labelled by phalloidin-FITC (#P5282, Merck). UdRPC were also subjected to flow cytometry using the CyAn ADP (Beckman Coulter, Indianapolis, IA, USA) to specifically detect renal progenitor cell surface markers by using antibodies directed against CD24-FITC (#FCMAB188F, Merck), VCAM- 1/CD106-PE (#FAB5649P, R&D Systems), and CD133-APC (#FAB11331A, R&D Systems).

### Differentiation of UdRPC into podocytes and tubular cells

Podocytes differentiation was initiated after culturing UdRPC on rat collagen type I coated plates to a confluency of approximately 60-70% in advanced RPMI medium supplemented with 30 μM retinoic acid for 14 days ^10^. Expression of α-actinin 4 (#AB108198, Abcam, Cambridge, UK), podocin (NPHS2, #AB50339, Abcam), nephrin (NPHS1 adhesion molecule, #AB4626861, Thermo Fisher Scientific), synaptopodin (SYNPO, #A5-56997, Thermo Fisher Scientific) and the cytoskeletal marker phalloidin-Alexa Fluor 488 (#A12379, Thermo Fisher Scientific) was confirmed as described ^10^.

Tubular cell differentiation was initiated after culturing undifferentiated UdRPC on laminin-421-coated plates to a confluency of approximately 90% (laminin-421, #LN421-02, BioLamina, Sundbyberg, Sweden). The cells were induced to differentiate in renal epithelial cell growth medium (REGM) supplemented with 10 ng/ml bone morphogenetic protein 2 (BMP2) and 2.5 ng/ml BMP7 for seven days ^35^. The tubular-associated markers zonular occludens-1 (ZO-1, #33-9100, Thermo Fisher Scientific), Na^+^/K^+^-ATPase (#ab76020, Abcam), chloride voltage-gated channel Kb (CLCNKB, #AB236733, Abcam), and SRY-box transcription factor 17 (SOX17, #AF1924, R&D) were confirmed by immunofluorescence.

### UdRPC transplantation

The animal experiments were performed by Phenos GmbH (Hannover, Germany) according to our specifications. All procedures were approved by the local authorities for animal protection (Niedersächsisches Landesamt für Verbraucherschutz und Lebensmittelsicherheit; reference number 33.19-42502-04-22-00107) and were performed in accordance with the German animal welfare act and the ARRIVE guidelines. A total of 21 male C57Bl/6N mice (12-15 weeks old; Charles River, Sulzfeld, Germany) were used for the present study (supplemental Table S1, S2). The animals received a standard diet with free access to food and water in an enriched environment and were monitored daily. To ensure analgesia, Buprenorphine and Metamizole were administered subcutaneously prior to surgery. As a contralateral control (control I; n = 21), 20 µl phosphate buffered saline (PBS) were injected under the renal capsule of the right kidney of all animals used for the study. In the sham group (group 1; control II; n = 5), 20 µl of UdRPC suspension (1x10^6^ cells) were injected under the renal capsule of the left kidney. In group 2 and 3 the left renal pedicle was clamped with a microaneurysm clip for 45 min (IRI model). In group 2 (n = 8), 20 µl of the vehicle PBS were injected into the left kidney. The group 3 (n = 8) received also 1x10^6^ UdRPC in 20 µl PBS under the renal capsule of the clamped left kidney. Body weight was measured, and blood samples were taken prior to surgery (day 0) and on day 1, 7, 14, and 21. On day 21 the mice were euthanized in deep isoflurane anesthesia by blood sampling via the left ventricle. After terminal blood collection whole body perfusion was performed with ice cold PBS via the left ventricle which results in circulatory arrest. The fresh weight of the kidneys was subsequently determined.

### Clinical chemistry and proteome analyses

For assessing renal function, s-creatinine and blood urea nitrogen (BUN) were measured in serum samples of mice on day 0, 1, 7, 14, and 21 using the Olympus analyzer (AU400) by the Phenos GmbH. Serum samples from mice were further analyzed on day 0, 14, and 21 using the Proteome Profiler Human Kidney Biomarker Array Kit (ARY019; R&D Systems, Minneapolis, MN, USA) according to the manufacturer’s recommendations. Collected blood samples were pooled from distinct animals based on their experimental group assignment to obtain sufficient amounts of protein. The secretome of UdRPC UF31 was analyzed by the Proteome Profiler Human XL Cytokine Array Kit (ARY022B; R&D Systems) in cell culture medium as specified by the manufacturer. Image analysis including grid-finding and quantification was semi-automatically performed using the FIJI/ImageJ software ^36^, normalized via the mean of the reference spots in the R/Bioconductor ^37^ and assessed for gene expression via detection-p-values < 0.05 as described ^38^. GO over-representation analysis was performed via the R package GOstats ^39^ using the total list of proteins of the Proteome Profiler Human XL Cytokine Array as background. For the Venn diagram comparison of proteins detected in the secretome of UdRPC UF31 and in animal blood samples only the overlap of cytokines between both array types ARY019 and ARY022B was considered.

### Sirius Red staining

The amount of extracellular matrix protein (ECM) was quantified after Sirius Red staining of paraffin-embedded mouse kidney sections (10 µm). The Pico-Sirius Red Stain Kit (#SCY-PSR-1, Biozol Diagnostica GmbH, Echingen, Germany) was used according to the protocol provided by the manufacturer. Stained areas of the kidney sections were quantified using the CellSens Dimension software (version 1.16, Evident Europe GmbH, Hamburg, Germany).

### Reverse transcriptase quantitative polymerase chain reaction (RT-qPCR)

TaqMan Reverse Transcription Reagents (#N8080234, Applied Biosystems) were used to synthesize cDNA from 1 µg total RNA per sample. For the PCR, 10 ng cDNA, 0.6 µM from each primer set (supplemental Table S3), and the Takyon No Rox SYBR Green MasterMix dTTP Blue (#UF-NSMT-B0701, Eurogentec, Seraing, Belgium) were mixed. The PCR products were amplified by applying a standard protocol. The CT values were analyzed by the 2(−ΔΔCt) method and normalized against the housekeeping gene hypoxanthine-guanine-phosphoribosyl transferase 1 (*Hprt1*).

### In-situ hybridization of ALU repeats

To identify transplanted UdRPC, in-situ hybridization (ISH) of human-specific ALU repeat sequences was performed on paraffin-embedded mouse kidney sections after target retrieval through Tris-EDTA buffer (10 mM Tris base, 1 mM EDTA solution, 0.05% Tween 20, pH 9.0) treatment for 30 min at 85°C and blocking of peroxidase activity (Dual Enzyme block; #S2003, Dako-Agilent, Santa Clara, CA, USA). The tissue sections were permeabilized using proteinase K (10 µg/ml, #P2308, Merck) diluted in digestion buffer (2x saline sodium citrate buffer/SSC, 2.5% sodium dodecyl sulfate). The ZytoFast DNA (+) control probe with digoxigenin (#T-1053-100; ZytoVision GmbH, Bremerhaven, Germany) was directly layered onto each section, denatured at 75°C for 5 min, and hybridized at 37°C for 2 hours. DIGX rabbit anti-digoxigenin antibody (#ENZ-ABS303; Enzo Biochem Inc., Long Island, NY, USA), anti-rabbit antibody coupled with horseradish peroxidase/HRP (#K4010, Dako-Agilent), and 3,3’-diaminobenzidine (SIGMAFAST DAB, #D4293, Merck) was applied to stain the bound probes. The cell nuclei were counterstained by diluted hematoxylin solution (#1.05175.0500, Merck).

### Statistics

Statistical analyses were performed using GraphPad Prism software version 10.4.1 (GraphPad Software, San Diego, CA, USA). Statistical significance was tested by one way analysis of variance (ANOVA). Significant differences were indicated by asterisks. Results are presented in scatter plots and the arithmetic mean as well as standard error of mean (SEM) is indicated.

## Supporting information

Supplement - Human Urine-derived SIX2-positive renal progenitor cells improve kidney injury in an IRI mouse model

## List of supplementary materials

Supplemental Figure S1: Proteins expressed on the Proteome Profiler Human Kidney Biomarker Array Kit (ARY019) are associated with inflammation and vascular processes.

Supplemental Figure S2: Membranes of the Proteome Profiler Human Kidney Biomarker Array Kit (ARY019) used for this study to analyze serum samples.

Supplemental Table S1: List of mice used for the experiments and labelling of tissue sections.

Supplemental Table S2: Body weight of mice before (day 0/d0) and after surgery.

Supplemental Table S3: Mouse primer sets for RT-qPCR.

Supplemental Table S4: GO term analysis of cytokines released by cultured UdRPC UF31.

### Acknowledgements

The authors are grateful to the Phenos GmbH (Hannover, Germany) for performing the in vivo experiments.

## Author contributions

CK, LS, JA: wrote the manuscript; LS, AN, MB: isolated and maintained UdRPC; LS, AN, CT, JA: characterization of UdRPC; LS, AN: transplantation management; WW: carried out bioinformatics; CK: histological examination and microscopy; CK, MB: performed RT-PCR analyses; CT, LE: conducted secretome analyses. All authors approved the manuscript.

## Data and materials availability

The authors declare that all data supporting the findings of this study are available within the article and its supplementary material.

## REFERENCES

1. Basile, D. P., Anderson, M. D. & Sutton, T. A. Pathophysiology of Acute Kidney Injury. Comprehensive Physiology 2, 1303–1353 (2012).

2. Hoste, E. A. J. et al. Global epidemiology and outcomes of acute kidney injury. Nature Reviews Nephrology 14, 607–625 (2018).

3. Kurzhagen, J. T., Dellepiane, S., Cantaluppi, V. & Rabb, H. AKI: an increasingly recognized risk factor for CKD development and progression. Journal of Nephrology 33, 1171–1187 (2020).

4. Sundström, J. et al. Prevalence, outcomes, and cost of chronic kidney disease in a contemporary population of 2·4 million patients from 11 countries: The CaReMe CKD study. Lancet 20, 100438 (2022).

5. Ortiz, A., Mattace-Raso, F., Soler, M. J. & Fouque, D. Ageing meets kidney disease. Nephrology Dialysis Transplantation 38, 523–526 (2023).

6. Francis, A. et al. Chronic kidney disease and the global public health agenda: an international consensus. Nature Reviews Nephrology 20, 473–458 (2024).

7. Yu, P. et al. Beyond waste: understanding urine’s potential in precision medicine. Trends in Biotechnology 42, 953–969 (2024).

8. Bohndorf, M. et al. Derivation and characterization of integration-free iPSC line ISRM-UM51 derived from SIX2-positive renal cells isolated from urine of an African male expressing the CYP2D6 *4/*17 variant which confers intermediate drug metabolizing activity. Stem Cell Research 25, 18–21 (2017).

9. Rahman, M. S., et al. The FGF, TGFβ and WNT axis Modulate Self-renewal of Human SIX2+ Urine Derived Renal Progenitor Cells. Scientific Reports 10, 739 (2020).

10. Erichsen, L. et al. Activation of the Renin–Angiotensin System Disrupts the Cytoskeletal Architecture of Human Urine-Derived Podocytes. Cells 11, 1095 (2022).

11. Kobayashi, A. et al. Six2 Defines and Regulates a Multipotent Self-Renewing Nephron Progenitor Population throughout Mammalian Kidney Development. Cell Stem Cell 3, 169–181 (2008).

12. Dong, X. et al. Beneficial effects of urine-derived stem cells on fibrosis and apoptosis of myocardial, glomerular and bladder cells. Molecular and Cellular Endocrinology 427, 21–32 (2016).

13. Tian, S. F. et al. Human urine-derived stem cells contribute to the repair of ischemic Acute kidney injury in rats. Molecular Medicine Reports 16, 5541–5548 (2017).

14. Li, X. et al. Human urine-derived stem cells protect against renal ischemia/reperfusion injury in a rat model via exosomal miR-146a-5p which targets IRAK1. Theranostics 10, 9561–9578 (2020).

15. Zhang, C. et al. Reno-protection of urine-derived stem cells in a chronic kidney disease rat model induced by renal ischemia and nephrotoxicity. International Journal of Biological Sciences 16, 435–446 (2020).

16. Gao, W. W. et al. Locally transplanted human urine-induced nephron progenitor cells contribute to renal repair in mice kidney with diabetic nephropathy. Biochemical and Biophysical Research Communications 629, 128–134 (2022).

17. Bonventre, J. v. & Yang, L. Cellular pathophysiology of ischemic acute kidney injury. Journal of Clinical Investigation 121, 4210–4221 (2011).

18. Hueper, K. et al. Acute Kidney Injury: Arterial Spin Labeling to Monitor Renal Perfusion Impairment in Mice—Comparison with Histopathologic Results and Renal Function. Radiology 270, 117–124 (2014).

19. Thorenz, A. et al. IL-17A blockade or deficiency does not affect progressive renal fibrosis following renal ischaemia reperfusion injury in mice. Journal of Pharmacy and Pharmacology 69, 1125–1135 (2017).

20. Colantuoni, M. et al. Constitutive IL-1RA production by modified immune cells protects against IL-1–mediated inflammatory disorders. Science Translational Medicine 15, eade3856 (2023).

21. Saxton, R. A. et al. Structure-based decoupling of the pro- and anti-inflammatory functions of interleukin-10. Science 371, eabc8433 (2021).

22. Suzuki, S. et al. Clinicopathological significance of platelet-derived growth factor (PDGF)-B and vascular endothelial growth factor-A expression, PDGF receptor-β phosphorylation, and microvessel density in gastric cancer. BMC Cancer 10, 659 (2010).

23. Lin, Y. et al. Role of endothelial PDGFB in arterio-venous malformations pathogenesis. Angiogenesis 27, 193–209 (2024).

24. Cressman, A. et al. Investigational New Drug-enabling studies to use genetically modified mesenchymal stromal cells in patients with critical limb ischemia. Stem Cells Translational Medicine 14, szae094 (2025).

25. Slater, T., Haywood, N. J., Matthews, C., Cheema, H. & Wheatcroft, S. B. Insulin-like growth factor binding proteins and angiogenesis: from cancer to cardiovascular disease. Cytokine and Growth Factor Reviews 46, 28–35 (2019).

26. Li, T. et al. IGFBP2: integrative hub of developmental and oncogenic signaling network. Oncogene 39, 2243–2257 (2020).

27. Wang, S. et al. Growth differentiation factor 15 promotes blood vessel growth by stimulating cell cycle progression in repair of critical-sized calvarial defect. Scientific Reports 7, 9027 (2017).

28. Ha, G. et al. GDF15 secreted by senescent endothelial cells improves vascular progenitor cell functions. PLOS ONE 14, e0216602 (2019).

29. Ren, Z. et al. Contribution of alterations in peritubular capillary density and microcirculation to the progression of tubular injury and kidney fibrosis. The Journal of Pathology 266, 95–108 (2025).

30. Moore, B. B. et al. GM-CSF Regulates Bleomycin-Induced Pulmonary Fibrosis Via a Prostaglandin-Dependent Mechanism. The Journal of Immunology 165, 4032–4039 (2000).

31. Liu, L. et al. Sympathetic nerve promotes renal fibrosis by activating M2 macrophages through β2-AR-Gsa. Clinical Immunology 270, 110397 (2025).

32. Hu, B. et al. Sensory nerves regulate mesenchymal stromal cell lineage commitment by tuning sympathetic tones. Journal of Clinical Investigation 130, 3483–3498 (2020).

33. Cohen, T., Nahari, D., Cerem, L. W., Neufeld, G. & Levi, B.-Z. Interleukin 6 Induces the Expression of Vascular Endothelial Growth Factor. Journal of Biological Chemistry 271, 736–741 (1996).

34. Wang, M.-J. et al. The double-edged effects of IL-6 in liver regeneration, aging, inflammation, and diseases. Experimental Hematology & Oncology 13, 62 (2024).

35. Mboni-Johnston, I. M. et al. Sensitivity of Human Induced Pluripotent Stem Cells and Thereof Differentiated Kidney Proximal Tubular Cells towards Selected Nephrotoxins. International Journal of Molecular Sciences 25, 81 (2023).

36. Schneider, C. A., Rasband, W. S. & Eliceiri, K. W. NIH Image to ImageJ: 25 years of image analysis. Nature Methods 9, 671–675 (2012).

37. Gentleman, R. C. et al. Bioconductor: open software development for computational biology and bioinformatics. Genome Biology 5, R80 (2004).

38. Wruck, W. et al. Urine-Based Detection of Biomarkers Indicative of Chronic Kidney Disease in a Patient Cohort from Ghana. Journal of Personalized Medicine 13, 38 (2022).

39. Falcon, S. & Gentleman, R. Using GOstats to test gene lists for GO term association. Bioinformatics 23, 257–258 (2007).

