## Supplement - Human Urine-derived SIX2-positive renal progenitor cells improve kidney injury in an IRI mouse model for "Human Urine-derived SIX2-positive renal progenitor cells improve kidney injury in an IRI mouse model"

### SUPPLEMENTAL TABLES AND FIGURES

***Supplemental Table S1: List of mice used for the experiments and labelling of tissue sections.***

| Group | Surgery | Treatment injured left kidney | Treatment contralateral right kidney | Animal | Date of surgery | Date of sacrifice | Left kidney FFPE sampels | Contralateral right kidney FFPE sampels |
|---|---|---|---|---|---|---|---|---|
| 1 | Sham | UdRPC | Vehicle PBS | 1 | 13.09.2022 | 04.10.2022 | PH 2022-58 A | PH 2022-58 B |
| | | | | 22 | 13.09.2022 | 04.10.2022 | PH 2022-59 A | PH 2022-59 B |
| | | | | 3 | 13.09.2022 | 04.10.2022 | PH 2022-60 A | PH 2022-60 B |
| | | | | 23 | 13.09.2022 | 04.10.2022 | PH 2022-61 A | PH 2022-61 B |
| | | | | 21 | 13.09.2022 | 04.10.2022 | PH 2022-62 A | PH 2022-62 B |
| 2 | IRI | Vehicle PBS | Vehicle PBS | 5 | 13.09.2022 | 04.10.2022 | PH 2022-63 A | PH 2022-63 B |
| | | | | 6 | 13.09.2022 | 04.10.2022 | PH 2022-64 A | PH 2022-64 B |
| | | | | 24 | 22.09.2022 | 13.10.2022 | PH 2022-65 A | PH 2022-65 B |
| | | | | 25 | 22.09.2022 | 13.10.2022 | PH 2022-66 A | PH 2022-66 B |
| | | | | 9 | 13.09.2022 | 04.10.2022 | PH 2022-67 A | PH 2022-67 B |
| | | | | 10 | 13.09.2022 | 04.10.2022 | PH 2022-68 A | PH 2022-68 B |
| | | | | 11 | 13.09.2022 | 04.10.2022 | PH 2022-69 A | PH 2022-69 B |
| | | | | 12 | 13.09.2022 | 04.10.2022 | PH 2022-70 A | PH 2022-70 B |
| 3 | IRI | UdRPC | Vehicle PBS | 13 | 22.09.2022 | 13.10.2022 | PH 2022-71 A | PH 2022-71 B |
| | | | | 14 | 22.09.2022 | 13.10.2022 | PH 2022-72 A | PH 2022-72 B |
| | | | | 15 | 22.09.2022 | 13.10.2022 | PH 2022-73 A | PH 2022-73 B |
| | | | | 16 | 22.09.2022 | 13.10.2022 | PH 2022-74 A | PH 2022-74 B |
| | | | | 17 | 22.09.2022 | 13.10.2022 | PH 2022-75 A | PH 2022-75 B |
| | | | | 18 | 22.09.2022 | 13.10.2022 | PH 2022-76 A | PH 2022-76 B |
| | | | | 19 | 22.09.2022 | 13.10.2022 | PH 2022-77 A | PH 2022-77 B |
| | | | | 20 | 22.09.2022 | 13.10.2022 | PH 2022-78 A | PH 2022-78 B |

### *Supplemental Table S2: Body weight of mice before (day 0/d0) and after surgery.*

| Group | Surgery | Treatment injured left kidney | Treatment contralateral right kidney | Animal | Body weight d0 [g] | Body weight d1 [g] | Body weight d2 [g] | Body weight d3 [g] | Body weight d7 [g] | Body weight d14 [g] | Body weight d21 [g] |
|---|---|---|---|---|---|---|---|---|---|---|---|
| 1 | Sham | UdRPC | Vehicle PBS | 1 | 24.0 | 22.3 | 23.0 | 23.8 | 24.2 | 24.3 | 25.0 |
| | | | | 22 | 25.4 | 25.9 | 26.3 | 26.5 | 25.0 | 27.1 | 26.5 |
| | | | | 3 | 24.3 | 24.4 | 24.9 | 25.4 | 25.2 | 26.4 | 27.1 |
| | | | | 23 | 24.8 | 22.7 | 23.1 | 24,3 | 24.1 | 24.5 | 24.4 |
| | | | | 21 | 27.9 | 26.4 | 26.5 | 26.7 | 25.9 | 26.7 | 26.3 |
| 2 | IRI | Vehicle PBS | Vehicle PBS | 5 | 25.6 | 23.0 | 23.8 | 23.9 | 24.2 | 25.0 | 24.9 |
| | | | | 6 | 22.5 | 19.5 | 19.6 | 20.5 | 22.0 | 23.4 | 21.8 |
| | | | | 24 | 25.7 | 24.2 | 24.6 | 24.5 | 24.7 | 24.9 | 24.6 |
| | | | | 25 | 27.0 | 24.6 | 24.7 | 24.9 | 25.0 | 26.3 | 27.1 |
| | | | | 9 | 24.4 | 22.5 | 22.8 | 23.4 | 23.8 | 23.4 | 23.4 |
| | | | | 10 | 24.9 | 18.6 | 19.6 | 19.8 | 17.4 | 18.5 | 18.6 |
| | | | | 11 | 20.3 | 23.4 | 23.9 | 24.2 | 24.2 | 24.0 | 23.9 |
| | | | | 12 | 24.3 | 23.1 | 23.6 | 23.7 | 24.2 | 24.6 | 24.2 |
| 3 | IRI | UdRPC | Vehicle PBS | 13 | 21.3 | 19.8 | 20.0 | 20.1 | 21.3 | 21.1 | 21.7 |
| | | | | 14 | 25.1 | 23.4 | 23.4 | 23.3 | 24.7 | 25.5 | 25.1 |
| | | | | 15 | 22.6 | 21.9 | 22.3 | 22.5 | 22.3 | 23.6 | 23.1 |
| | | | | 16 | 24.8 | 23.4 | 23.3 | 23.4 | 24.2 | 23.9 | 23.4 |
| | | | | 17 | 26.7 | 24.0 | 24.5 | 24.6 | 27.0 | 26.4 | 26.2 |
| | | | | 18 | 24.9 | 24.1 | 24.3 | 24.4 | 25.3 | 27.0 | 27.0 |
| | | | | 19 | 23.6 | 22.4 | 22.5 | 22.7 | 24.1 | 24.8 | 25.2 |
| | | | | 20 | 20.3 | 18.6 | 18.8 | 19.2 | 20.6 | 21.4 | 20.7 |

### *Supplemental Table S3: Mouse primer sets for RT-qPCR.*

| Gene | Forward Primer | Reverse Primer | Accession No. |
|---|---|---|---|
| ***Ccl2 (Mcp-1)*** | CCACTCACCTGCTGCTACTC | CTTCTTGGGGTCAGCACAGA | NM_011333.3 |
| ***Col1α2*** | AGTTTCATCTGGCCCTGGAC | TCCAACGACTCCTCTCTCCC | NM_007743.3 |
| ***Col3α1*** | AATTGGGATGCAGCCACCTT | TTGAGGTCCATGGCCATCAG | NM_009930.2 |
| ***Col4α1*** | CACAAAAGGGACGAGGGGAC | GGCCGAGAATTTCACCAGGA | NM_009931.2 |
| ***Ctgf (Ccn2)*** | AGCGGTGAGTCCTTCCAAAG | TGGGCCAAATGTGTCTTCCA | NM_010217.2 |
| ***Hprt1*** | CAAGCTTGCTGGTGAAAAGGA | TCCAGTTTCACTAATGACACA | NM_013556.2 |
| ***Icam1*** | TTCTCATGCCGCACAGAACT | CGAGCTTCAGAGGCAGGAAA | NM_010493.3 |
| ***Sox9*** | CCAGCAAGAACAAGCCACAC | TCGGGGTGGTCTTTCTTGTG | NM_011448.4 |

**A** Proteome Profiler Human Kidney Biomarker Array Kit (ARY019, serum samples on day 0)

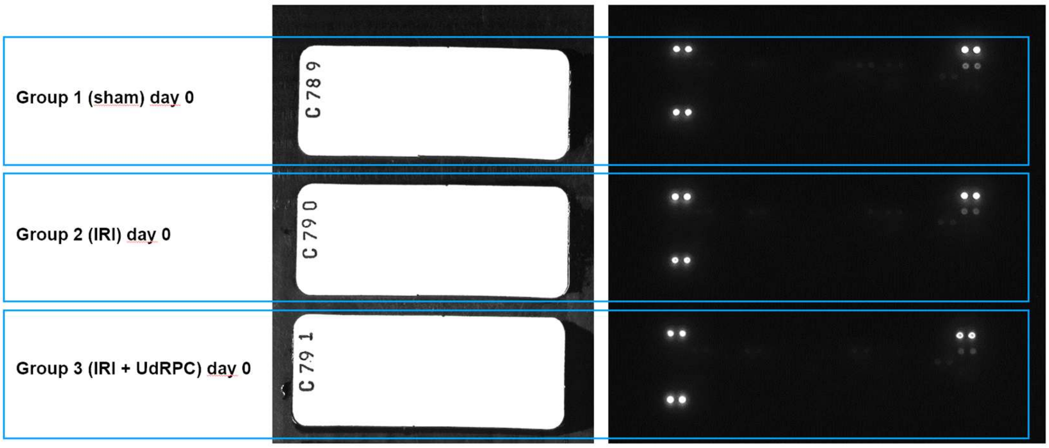


**B** Proteome Profiler Human Kidney Biomarker Array Kit (ARY019, serum samples on day 14)

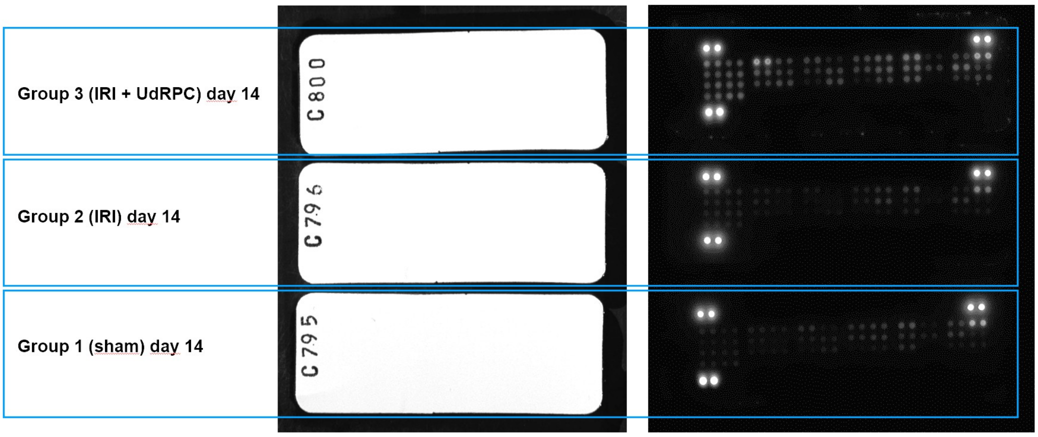


**C** Proteome Profiler Human Kidney Biomarker Array Kit (ARY019, serum samples on day 21)

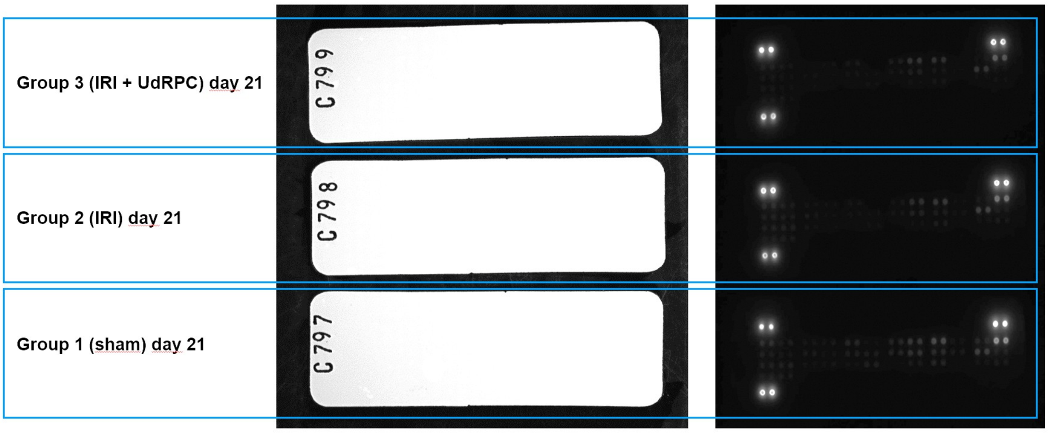


***Supplemental Figure S1: Membranes of the Proteome Profiler Human Kidney Biomarker Array Kit (ARY019) used for this study to analyze serum samples.*** *Membranes of the 3 experimental groups for serum samples on (**A**) day 0, (**B**) day 14, and (**C**) day 21.*

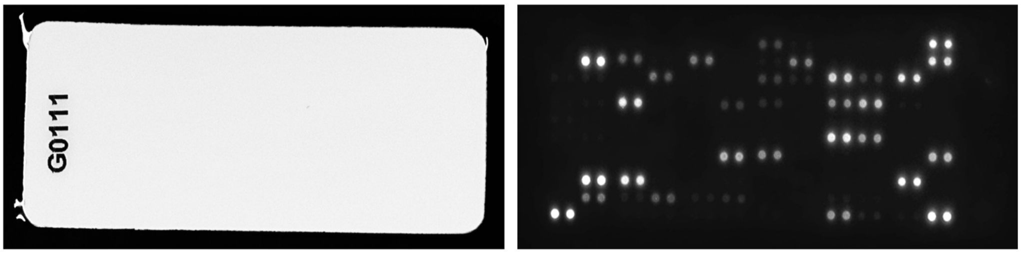


***Supplemental Figure S2: The secretome of UdRPC UF31 was analyzed by the Proteome Profiler Human XL Cytokine Array Kit (ARY022B; R&D Systems) in cell culture medium.***

***Supplemental Table S4: GO term analysis of cytokines released by cultured UdRPC UF31.*** *The table indicates GO terms with reference to biological processes (BP) and cellular components (CC).*

| GO ID | Pvalue | Odds Ratio | Exp. Count | Count | Size | Term | GO Type | Genes |
|---|---|---|---|---|---|---|---|---|
| GO:0048259 | 0.02 | 9.3 | 3.7 | 7 | 8 | regulation of receptor-mediated endocytosis | BP | ADIPOQ,CCL19,DKK1,EGF,IL4,SERPINE1,VEGFA |
| GO:0030168 | 0.02 | ∞ | 2.3 | 5 | 5 | platelet activation | BP | CD40LG,IL6,PDGFA,PDGFB,PF4 |
| GO:0030195 | 0.02 | ∞ | 2.3 | 5 | 5 | negative regulation of blood coagulation | BP | PDGFA,PDGFB,PLAUR,SERPINE1,THBS1 |
| GO:0046718 | 0.02 | ∞ | 2.3 | 5 | 5 | symbiont entry into host cell | BP | BSG,DPP4,ICAM1,PTX3,TFRC |
| GO:0050818 | 0.02 | ∞ | 2.3 | 5 | 5 | regulation of coagulation | BP | PDGFA,PDGFB,PLAUR,SERPINE1,THBS1 |
| GO:0051701 | 0.02 | ∞ | 2.3 | 5 | 5 | biological process involved in interaction with host | BP | BSG,DPP4,ICAM1,PTX3,TFRC |
| GO:1900046 | 0.02 | ∞ | 2.3 | 5 | 5 | regulation of hemostasis | BP | PDGFA,PDGFB,PLAUR,SERPINE1,THBS1 |
| GO:0071840 | 0.02 | 2.5 | 25.6 | 31 | 55 | cellular component organization or biogenesis | BP | ADIPOQ,ANG,APOA1,CCL19,CCL2,CSF2,CST3,CXCL1,CXCL12,DKK1,DPP4,EGF,GDF15,ICAM1,IGFBP3,IL17A,IL4,IL6,KLK4,LIF,MIF,MPO,PDGFA,PDGFB,PLAUR,PTX3,SERPINE1,SPP1,TFRC,THBS1,VEGFA |
| GO:0001894 | 0.03 | 5.4 | 4.6 | 8 | 10 | tissue homeostasis | BP | CSF1,IL17A,IL6,RBP4,SPP1,TFF3,TFRC,VEGFA |
| GO:0050878 | 0.03 | ∞ | 1.7 | 4 | 4 | regulation of body fluid levels | BP | CD40LG,IL6,PDGFA,PDGFB,PF4,PLAUR,SERPINE1,THBS1,VEGFA |
| GO:0042060 | 0.03 | 4.1 | 5.6 | 9 | 12 | wound healing | BP | CD40LG,IL6,PDGFA,PDGFB,PF4,PLAUR,SERPINE1,THBS1,VEGFA |
| GO:0015850 | 0.04 | 7.8 | 3.3 | 6 | 7 | organic hydroxy compound transport | BP | ADIPOQ,APOA1,CXCL12,EGF,RBP4,SPP1 |
| GO:0120035 | 0.04 | 7.8 | 3.3 | 6 | 7 | regulation of plasma membrane bounded cell projection organization | BP | CCL19,CXCL12,DKK1,ICAM1,SPP1,VEGFA |
| GO:0048519 | 0,04 | 2.5 | 32.5 | 37 | 70 | negative regulation of biological process | BP | ADIPOQ,ANG,APOA1,CCL19,CCL2,CD40LG,CSF2,CST3,CXCL1,CXCL12,DKK1,DPP4,EGF,GDF15,HAVCR2,ICAM1,IGFBP2,IGFBP3,IL17A,IL24,IL4,IL6,LIF,MIF,MPO,PDGFA,PDGFB,PF4,PLAUR,PTX3,RBP4,RETN,SERPINE1,SPP1,TFRC,THBS1,VEGFA |
| GO:0051129 | 0.04 | 3.4 | 6.5 | 10 | 14 | negative regulation of cellular component organization | BP | ADIPOQ,APOA1,CST3,DKK1,DPP4,LIF,SPP1,TFRC,THBS1,VEGFA |
| GO:0001656 | 0.04 | ∞ | 1.9 | 4 | 4 | metanephros development | BP | ADIPOQ,LIF,PDGFA,PDGFB |
| GO:0001952 | 0.04 | ∞ | 1.9 | 4 | 4 | regulation of cell-matrix adhesion | BP | CSF1,SERPINE1,THBS1,VEGFA |
| GO:0010811 | 0.04 | ∞ | 1.9 | 4 | 4 | positive regulation of cell-substrate adhesion | BP | APOA1,CSF1,PDGFB,VEGFA |
| GO:0016485 | 0.04 | ∞ | 1.9 | 4 | 4 | protein processing | BP | DPP4,PLAUR,SERPINE1,THBS1 |
| GO:0031952 | 0.04 | ∞ | 1.9 | 4 | 4 | regulation of protein autophosphorylation | BP | ADIPOQ,PDGFA,PDGFB,VEGFA |
| GO:0042475 | 0.04 | ∞ | 1.9 | 4 | 4 | odontogenesis of dentin-containing tooth | BP | BSG,CSF1,KLK4,SERPINE1 |
| GO:0045017 | 0.04 | ∞ | 1.9 | 4 | 4 | glycerolipid biosynthetic process | BP | ANG,APOA1,PDGFA,PDGFB |
| GO:0048008 | 0.04 | ∞ | 1.9 | 4 | 4 | platelet-derived growth factor receptor signaling pathway | BP | ADIPOQ,PDGFA,PDGFB,VEGFA |
| GO:0090407 | 0.04 | ∞ | 1.9 | 4 | 4 | organophosphate biosynthetic process | BP | APOA1,IL4,PDGFA,PDGFB |
| GO:1904037 | 0.04 | ∞ | 1.9 | 4 | 4 | positive regulation of epithelial cell apoptotic process | BP | CCL2,CD40LG,IL6,THBS1 |
| GO:0031983 | 0.0002 | 11.9 | 7.99 | 15 | 17 | vesicle lumen | CC | APOA1,CFD,CXCL1,EGF,LCN2,MIF,MPO,PDGFA,PDGFB,PF4,PTX3,RETN,SERPINE1,THBS1,VEGFA |
| GO:0012505 | 0.0004 | 4.5 | 21.2 | 30 | 45 | endomembrane system | CC | ADIPOQ,APOA1,BSG,CD40LG,CFD,CSF1,CST3,CXCL1,DKK1,EGF,GDF15,HAVCR2,IGFBP3,IL6,KLK4,LCN2,MIF,MPO,PDGFA,PDGFB,PF4,PLAUR,PTX3,RETN,SERPINE1,SPP1,TFF3,TFRC,THBS1,VEGFA |
| GO:0099503 | 0.001 | 4.9 | 12.7 | 20 | 27 | secretory vesicle | CC | APOA1,BSG,CFD,CST3,CXCL1,EGF,KLK4,LCN2,MIF,MPO,PDGFA,PDGFB,PF4,PLAUR,PTX3,RETN,SERPINE1,TFF3,THBS1,VEGFA |
| GO:0097708 | 0.005 | 3.2 | 18.3 | 25 | 39 | intracellular vesicle | CC | ANG,APOA1,BSG,CFD,CST3,CXCL1,DKK1,DPP4,EGF,HAVCR2,KLK4,LCN2,MIF,MPO,PDGFA,PDGFB,PF4,PLAUR,PTX3,RETN,SERPINE1,TFF3,TFRC,THBS1,VEGFA |
| GO:0065010 | 0.009 | 3.2 | 14.1 | 20 | 30 | extracellular membrane-bounded organelle | CC | APOA1,BSG,CFD,CST3,CXCL12,DPP4,EGF,GDF15,ICAM1,IGFBP2,LCN2,MIF,MPO,RBP4,RETN,SERPINE1,SHBG,SPP1,TFRC,THBS1 |
| GO:0070062 | 0.009 | 3.2 | 14.1 | 20 | 30 | extracellular exosome | CC | APOA1,BSG,CFD,CST3,CXCL12,DPP4,EGF,GDF15,ICAM1,IGFBP2,LCN2,MIF,MPO,RBP4,RETN,SERPINE1,SHBG,SPP1,TFRC,THBS1 |
| GO:0031093 | 0.009 | 10.7 | 4.2 | 8 | 9 | platelet alpha granule lumen | CC | CFD,EGF,PDGFA,PDGFB,PF4,SERPINE1,THBS1,VEGFA |
| GO:0034774 | 0.01 | 11.2 | 3.4 | 7 | 8 | secretory granule lumen | CC | APOA1,CFD,CXCL1,EGF,LCN2,MIF,MPO,PDGFA,PDGFB,PF4,PTX3,RETN,SERPINE1,THBS1,VEGFA |
| GO:0043229 | 0.02 | 2.6 | 27.3 | 33 | 58 | intracellular organelle | CC | ADIPOQ,ANG,APOA1,BSG,CD40LG,CFD,CSF1,CSF2,CST3,CXCL1,DKK1,DPP4,EGF,GDF15,HAVCR2,IGFBP3,IL6,KLK4,LCN2,MIF,MPO,PDGFA,PDGFB,PF4,PLAUR,PTX3,RETN,SERPINE1,SPP1,TFF3,TFRC,THBS1,VEGFA |
| GO:0005788 | 0.02 | 4.5 | 6.1 | 10 | 13 | endoplasmic reticulum lumen | CC | APOA1,CSF1,CST3,IGFBP3,IL6,PDGFA,PDGFB,PLAUR,SPP1,THBS1 |
| GO:0030312 | 0.03 | 3.3 | 7.99 | 12 | 17 | external encapsulating structure | CC | ADIPOQ,ANG,APOA1,CXCL12,GDF15,ICAM1,PDGFB,PF4,PTX3,SERPINE1,THBS1,VEGFA |
| GO:0043227 | 0.04 | 2.8 | 13.9 | 18 | 36 | membrane-bounded organelle | CC | ADIPOQ,ANG,APOA1,BSG,CD40LG,CFD,CSF1,CSF2,CST3,CXCL1,CXCL12,DKK1,DPP4,EGF,GDF15,HAVCR2,ICAM1,IGFBP2,IGFBP3,IL6,KLK4,LCN2,MIF,MPO,PDGFA,PDGFB,PF4,PLAUR,PTX3,RBP4,RETN,SERPINE1,SHBG,SPP1,TFF3,TFRC,THBS1,VEGFA |
| GO:0062023 | 0.05 | 3.3 | 6.6 | 10 | 14 | collagen-containing extracellular matrix | CC | ADIPOQ,ANG,APOA1,CXCL12,GDF15,ICAM1,PDGFB,PF4,SERPINE1,THBS1 |
| GO:0005769 | 0.05 | ∞ | 1.9 | 4 | 4 | early endosome | CC | APOA1,DKK1,HAVCR2,TFRC |
| GO:0036454 | 0.05 | ∞ | 1.9 | 4 | 4 | growth factor complex | CC | IGFBP3,PDGFA,PDGFB,VEGFA |
